# Single cell analysis of HIV expression and integration sites reveals robust viral expression across diverse chromatin environments

**DOI:** 10.1101/2025.06.19.660519

**Authors:** Yun Ma, Jackson J Peterson, David M Margolis, Edward P Browne

## Abstract

Entry of HIV into latency is determined by a combination of factors, including fluctuations in the viral Tat protein, as well as the transcriptomic phenotype of the host cell. Determining the impact of the proviral integration site on viral expression has been challenging due to difficulty in measuring integration site and viral expression from the same cell. To investigate the influence of the HIV integration site on HIV expression, we analyzed a combined scRNAseq/scATACseq dataset from 117,610 HIV infected primary CD4 T cells. We used the scATACseq data to recover HIV integration site information from 1530 cells, and correlated this information with viral RNA reads in the scRNAseq data. We observed that, overall, HIV expression did not differ depending on the genomic features of viral integration, such as genic vs non genic, intron vs exon and forward versus reverse orientation. Furthermore, we found that there was no significant difference in HIV expression across 15 distinct chromatin compartments. Additionally, analysis of expression for host genes that were the target of proviral integration revealed strong upregulation of expression of the targeted host gene in ∼5% of the infected cells. This insertional activation occurred almost exclusively when HIV was integrated in the same orientation as the host cell gene and occurred as a result of integration within diverse positions across a gene. These findings suggest that HIV expression is relatively robust to the genomic context of the HIV integration site, and that HIV can strongly upregulate expression of integration site genes during infection.

## Introduction

Since its introduction in the 1990s, antiretroviral therapy (ART) has transformed human immunodeficiency virus (HIV) from a fatal illness into a manageable chronic condition (Arts et al. 2012). ART blocks multiple steps of the viral life cycle, suppressing viral replication and reducing both disease burden and transmissibility. However, viral rebound typically occurs when ART is interrupted, indicating the persistence of a long-lived, replication-competent viral reservoir (Chun et al. 1997, Finzi et al. 1997, Siliciano et al. 2011). As a result, people with HIV (PWH) require lifelong therapy. This persistence is due to HIV latency, where integrated proviruses enter a transcriptionally silent state, evading immune detection and drugs targeting HIV (Mbonye et al. 2017). Like other retroviruses, HIV reverse-transcribes its single-stranded RNA genome into double-stranded DNA and integrates into the host genome (Craigie et al. 2012). A fraction of these proviruses become transcriptionally silent within infected cells, forming a latent reservoir that can reactivate and fuel rebound upon ART discontinuation (Chun et al. 2010, Yeh et al. 2021). With a half-life of approximately 44 months under ART, the latent reservoir poses a major barrier to viral eradication (Finzi et al. 1999, Siliciano et al. 2003). Thus, understanding the mechanisms that govern HIV latency is essential for developing HIV cure strategies.

HIV integrates into the host genome in a non-random fashion, favoring transcriptionally active regions (Schröder et al. 2002, Craigie et al. 2012). Integration places the provirus under the regulatory control of host chromatin, including histone modifications that influence gene expression. For example, H3K36me3 is associated with actively transcribed genes and mediates binding of LEDGF, a host factor that tethers HIV integrase to active chromatin (Craigie et al. 2012, Ernst et al. 2017). By contrast, marks like H3K9me3 are enriched in heterochromatin and associated with transcriptional silencing. Together, these epigenetic marks define local transcriptional environments that may influence HIV expression and latency.

HIV integration can also affect host gene regulation. Insertion into or near regulatory elements of host genes can result in abnormal transcription and clonal expansion of infected cells (Yeh et al. 2021). For instance, integration into the first intron of the STAT3 gene has been linked to overexpression and lymphoma development (Mellors et al. 2021). These cases highlight the potential for proviral integration to dysregulate host genes in a manner that contributes to reservoir persistence and pathogenesis. To better understand how integration site features shape both viral and host gene expression, we analyzed single-cell ATAC and RNA sequencing data from HIV-infected CD4+ T cells. Specifically, we examined HIV IS location (intron, exon, or intergenic), local chromatin states, and insertion orientation relative to host gene transcription. Our results confirm that HIV preferentially integrates into transcriptionally active genes and that orientation is largely random. Surprisingly, we found that HIV expression is relatively robust across a range of integration site environments, suggesting limited sensitivity to local chromatin context. However, we also observed that HIV frequently activates host gene expression when integrated in the same transcriptional direction, an orientation-dependent phenomenon with potential relevance to reservoir biology. Together, our findings shed new light on the interaction between HIV proviral integration and the host genome, with implications for understanding latency and reactivation.

## Results

### Identification of HIV integration sites from single cell ATACseq data

The influence of the genomic environment near HIV integration sites on HIV viral expression, including chromatin features and proviral transcriptional orientation, has not been fully examined. To investigate this, we leveraged a combined single cell RNAseq/ATACseq dataset that we had previously generated from a population of primary CD4+ T cells that were infected with a GFP-expressing reporter strain of HIV. Following infection, these cells were maintained in culture for two additional weeks while the cells returned to a resting state and downregulated HIV expression. At this time point, expression of HIV was variegated, with some cells exhibiting loss of HIV expression (latency), while other cells maintained active viral gene expression. Given that the ATACseq data contains reads that map to integrated proviruses, we speculated that these data would also include reads that contained HIV-human junctions points, thereby revealing the HIV integration sites in infected cells. To identify integration sites (IS) from these data, we aggregated scRNAseq/scATACseq data from eight HIV-infected CD4+ T cell samples we have previously reported (Manickam et al. 2024) and eight unpublished samples from a total of five different donors (**Table S1**). A subset of these samples had been stimulated with small molecules – either latency reversing agents (Vorinostat, Prostratin or iBET151) or the cannabinoid ι1-9-tetrahydrocannabinol (THC) for 24 hours prior to profiling. We then used the epiVIA pipeline established by Wang et al. (2020) to analyze paired-end ATAC sequencing data for HIV IS. In cases where the host-virus junction appeared in one sequencing read, the integration site could be directly identified. Otherwise, it was inferred when the two ends of a paired segment mapped to different organisms (termed “pair chimeric”, **Figure 1**).

**Figure 1.**
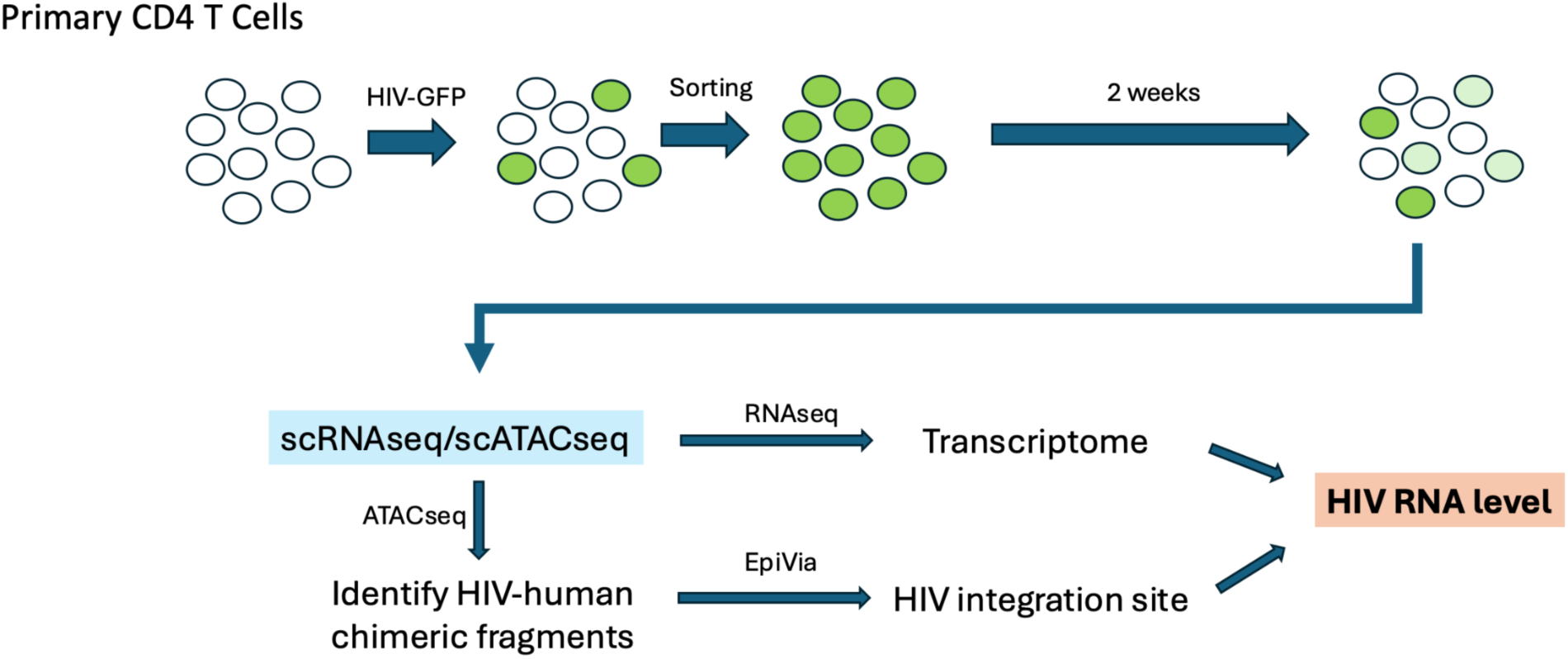
Schematic representation showing HIV infection, selection, and sequencing of single cells. Briefly, primary CD4 T cells from 5 different donors were activated then infected with a GFP expressing reporter strain of HIV. After two weeks of culture, a subset of cells were treated with small molecules for 24 hours, then the cells were profiled with a combined scRNAseq/scATACseq protocol. Chimeric HIV-human fragments were then identified from the ATACseq data using epiVIA. Cellular transcriptomes were also derived and used to determine viral RNA expression for each cell.

To validate that the identified HIV IS were not PCR recombination artifacts, we examined the position and orientation of the HIV-aligned ends in pair-chimeric reads relative to the viral genome. Notably, in ∼95% of chimeric fragments, the HIV-aligned reads localized within the viral long terminal repeats (LTRs) and were oriented toward the provirus termini (either 5′ or 3′). This non-random directional bias supports the conclusion that the captured HIV IS are unlikely to be PCR artifacts (**Figure S1A**). Additionally, the frequency of HIV–host chimeric fragments was significantly higher than that of baseline host–host chimeric fragments (**Figure S1B**). Initially, Chromosome 2 exhibited a significantly elevated recombination rate, primarily due to reads mapping to a long non-coding RNA LINC000486. This region was previously noted by Wang et al. (2020) and automatically excluded by their epiVIA pipeline. Upon removing reads aligned to this locus, chromosome 2’s recombination rate aligned with that of other host chromosomes. In total, we analyzed 117,610 single cells, of which approximately 40% contained ATACseq fragments that map to HIV and, among these, we identified HIV IS in 1,530 cells (1.3%). We hereafter refer to the subset of cells with identified HIV integration sites as the IS+ population.

### HIV expression distribution is influenced by treatment and batch effects

Some of the infected samples used in this study were exposed to small molecules before profiling (Vorinostat, Prostratin, iBET151 or THC), making treatment differences a potential confounding variable that could impact our results. For instance, Vorinostat (SAHA) and Prostratin are known latency reversal agents that can upregulate HIV expression (Manickam et al. 2024). Consistent with previous reports, we observed that HIV expression distributions in SAHA- and Prostratin-treated samples differed from those in DMSO controls (**Figure S2B**). In addition to treatment effects, batch effects are another important consideration, as single-cell expression matrices are highly sensitive to technical variation (Hafemeister et al. 2019). This is evident in **Figure S2A and S2B**, where unstimulated samples 1-2 exhibit a distinct expression distribution compared to unstimulated samples 3-5, in both total RNA and HIV expression. To address these sources of variation, we applied the Seurat integration pipeline to harmonize mRNA expression values across samples (Hao et al. 2023). After integration, expression distributions appeared more consistent and approximately normal across all samples (**Figure S2C**). To preserve the biological integrity of the data, we use the direct output of Seurat integration without additional transformations such as z-scoring or fold change. Because Seurat already corrects for sequencing depth and technical variation, further processing could introduce artificial scaling and obscure true biological signals (Hao et al. 2023). We therefore present the integrated expression distributions directly (**Figure S2D**). Overall, the integrated gene expression follows an approximately normal distribution with a right skew and a high density at zero, consistent with the sparse nature of single-cell data. For clarity, we will refer to the Seurat-integrated gene expression values as “Normalized gene expression” to distinguish between Seurat integration (which corrects for technical variation) and HIV integration (the biological process of HIV provirus incorporation into the host genome). These normalized values are unitless, as described above. For additional interpretability, we also include comparisons using unprocessed raw counts.

### IS+ cells exhibit a similar transcriptomic profile to IS- cells

To investigate whether the host transcriptome is associated with the recovery of an IS, we performed differential expression (DE)-based clustering on Seurat-integrated scRNAseq data. To ensure that variability in HIV expression did not drive clustering, we performed clustering on a version of the transcriptome with HIV expression excluded (**Figure 2A**). We found that the overall distribution of IS+ cells (N = 1,530) across transcriptional clusters closely mirrored that of the total cell population (N = 117,610), indicating that the IS+ cells represent a largely random sampling of the host cells at the transcriptomic level (**Figure 2B**). To assess whether the host cell transcriptional environment influences HIV expression, we first examined the distribution of treatment conditions across clusters to ensure that clustering was not confounded by treatments. The only exception to this was Cluster 4, which was enriched for Prostratin-treated cells, though this group comprised only 1.2% of the total population, and had no significant difference in HIV expression pattern (**Figure 2C**). Notably, Cluster 2 showed elevated total mRNA content (**Figure 2D**, **Figure S3A**). We confirmed the biological significance of this difference via ontology analysis of genes upregulated in this cluster, revealing enrichment for cell cycle related pathways, particularly those active during DNA synthesis and mitosis onset (**Figure S3C**). We also observed that HIV expression was elevated in this cluster regardless of whether cells were IS+ or IS- (**Figure 2E, Figure S3B, S3D**). In summary, DE-based clustering revealed that recovery of HIV integration sites from the data is not broadly associated with cellular gene expression patterns, and that HIV expression is influenced by the host cell transcriptional environment, consistent with previous reports (Bradley et al. 2018).

**Figure 2.**
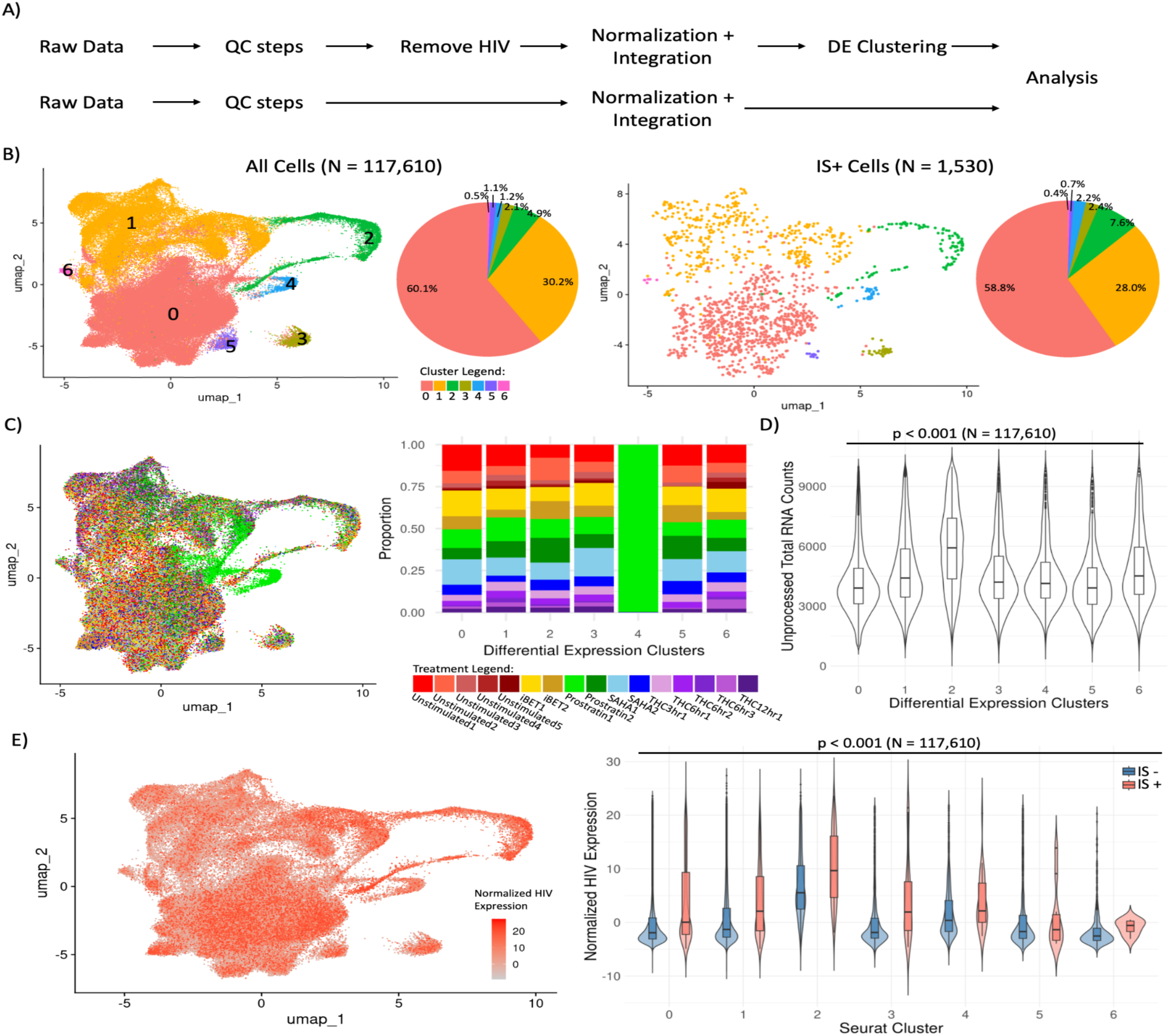
IS+ cells have a similar expression profile to IS- cells. **A)** Schematic of the data processing pipeline. Normalized HIV expression and clustering results were generated from independent copies of the raw single-cell RNA-seq data to prevent HIV transcripts from driving clustering. **B)** UMAP visualization of all cells (N = 117,610) and the IS+ subset (N = 1,530), colored by differential expression clusters. The accompanying pie chart shows the proportion of cells from each cluster within the population. **C)** UMAP plot of all cells colored by treatment condition. The bar plot shows the distribution of treatment conditions across DE-defined clusters. **D)** Combined boxplot and violin plot showing total mRNA content per cell across clusters. The Kruskal–Wallis test indicates significant differences across groups (p < 0.001, N = 117,610). **E)** UMAP visualization of normalized HIV expression in all cells. The accompanying boxplot and violin plot display the distribution of normalized HIV expression across DE clusters, stratified by IS status. Kruskal–Wallis tests show significant differences across clusters for both IS- and IS+ populations (p < 0.001 for each). Blue represents IS- cells; orange represents IS+ cells.

### IS+ cells are enriched for transcriptionally active proviruses

We next examined HIV expression and accessibility in the IS+ population compared to the overall infected cell population. Because HIV IS were sequenced using ATAC-seq, they are located within ATAC-accessible proviruses, which were open to host transcription factors and associated with elevated transcriptional activity. Based on this, we hypothesized that IS+ cells would exhibit higher levels of both HIV chromatin accessibility and RNA expression. Similar to what we have previously observed (Manickam et al. 2024), we observed a positive correlation between the number of HIV DNA fragments (estimates chromatin accessibility) and HIV RNA expression per cell (Spearman ρ = 0.41, **Figure 3A**). This trend was observed across both treated and untreated samples, indicating it was not driven by small-molecule treatment (ρ_untreated_ = 0.38, N_untreated_ = 31,082; ρ_treated_ = 0.41, N_treated_ = 86,528). When stratifying cells by HIV IS status, both IS+ and IS- cells followed the same correlation pattern between HIV DNA and RNA levels, despite a large difference in sample sizes (**Figure 3B**). As expected, no IS+ cells were found among those lacking HIV DNA fragments, since IS identification relies on the presence of HIV mapping DNA reads. Consistent with our hypothesis, IS+ cells contained significantly more HIV DNA fragments than IS- cells (**Figure 3C**). Additionally, IS+ cells exhibited higher HIV RNA expression compared to IS- cells (**Figure 3D**). These results confirm that the IS+ population is enriched for cells with transcriptionally active proviruses. Nevertheless, IS+ cells still displayed a wide range of expression levels from high to undetectable.

**Figure 3.**
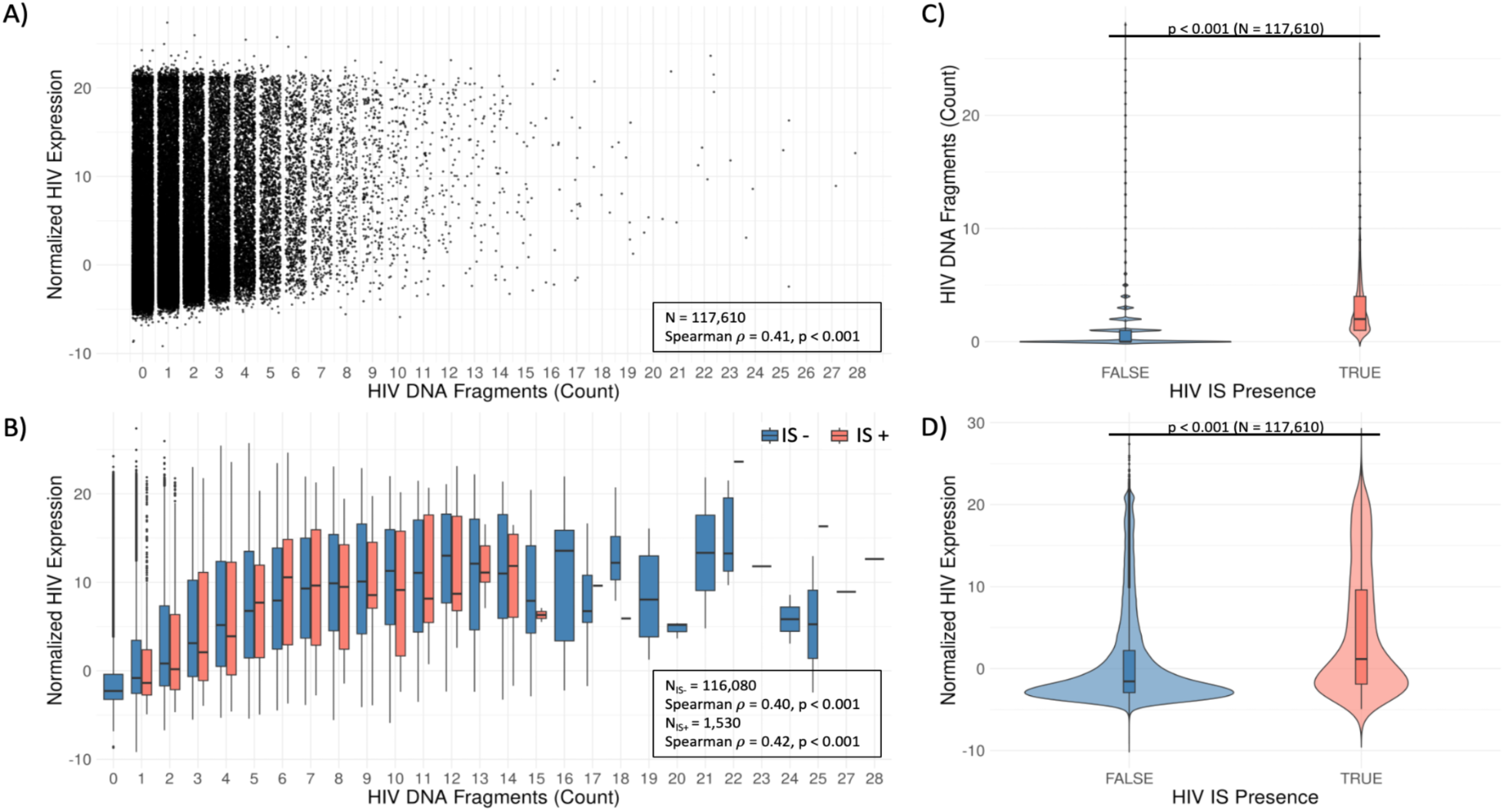
IS recovery is enriched for cells with ATAC-accessible and transcriptionally active proviruses. **A)** Scatter plot showing the correlation between HIV RNA expression and the number of HIV DNA fragments (from ATAC-seq) per cell. Each point represents a single cell. The Spearman correlation coefficient is ρ = 0.41 (p < 0.001, N=117,610). **B)** Box plot illustrating the same correlation as in panel A, stratified by HIV IS status. Blue represents IS- cells (ρ = 0.41, N=116,080), and orange represents IS+ cells (ρ = 0.43, N=1,530). **C)** Combined boxplot and violin plot showing the distribution of HIV DNA fragments per cell in IS+ (orange) and IS- (blue) populations. IS+ cells have significantly more HIV DNA fragments (Wilcoxon test, p < 0.001). **D)** Combined boxplot and violin plot showing the distribution of HIV RNA expression per cell in IS+ (orange) and IS- (blue) populations. IS+ cells have significantly higher HIV expression (Wilcoxon test, p < 0.001).

### HIV integration is enriched in transcriptionally active chromatin

Having identified integration sites in a population of HIV infected cells, we next investigated the genomic and epigenomic features surrounding HIV IS. Consistent with prior reports (Coffin et al. 2021), HIV IS were enriched in genic regions (**Figure 4A**). However, we found no significant difference in HIV expression between proviruses that were located within a gene (either in an exon or an intron) and those that were located in an intergenic region (p = 0.75, **Figure 4B**). This observation held true across treatment conditions and when examining raw counts (**Figure S4A, S4B**). To examine the chromatin configuration surrounding HIV IS at a finer resolution, we utilized chromHMM, a Hidden Markov Model-based annotation system developed by Roadmap Epigenomics Consortium et al. (2015). This model was trained on five histone marks (H3K4me3, H3K4me1, H3K36me3, H3K27me3, H3K9me3) at a 200 bp resolution across multiple cell types, including primary CD4^+^ T cells from peripheral blood (Roadmap Epigenomics Consortium et al. 2015). The chromHMM model assigns each 200 bp genomic region to one of 15 chromatin states based on histone mark emission densities, providing a reference for transcriptional and regulatory activity of the region (**Figure 4C**). We referenced this 15-state model as a benchmark for CD4^+^ T cell chromatin landscapes and annotated HIV IS accordingly. To compare our IS population to a previously published dataset of integration sites, we also examined 66,460 integration sites from HIV infected peripheral blood CD4 T cells (Coffin et al. 2021). We found that approximately 75% of HIV IS were located within transcriptionally active chromatin states, specifically states 4, 5, 6, and 7 (**Figure 4D**). This enrichment is significant, as those 4 states only account for 16.3% of the whole genome in base-pairs (**Figure 4C**). Furthermore, the chromatin state distribution of integration sites for our data strongly resembled the distribution from the reference set of integration sites (Coffin et al. 2021), indicating that the integration sites recovered from our IS+ cell population is likely representative of the overall population with respect to integration site pattern. In both our experimental dataset and the reference, approximately 20% of HIV IS were classified as “Quiescent.” These sites lacked the 5 core histone marks, making their chromatin context inconclusive.

**Figure 4.**
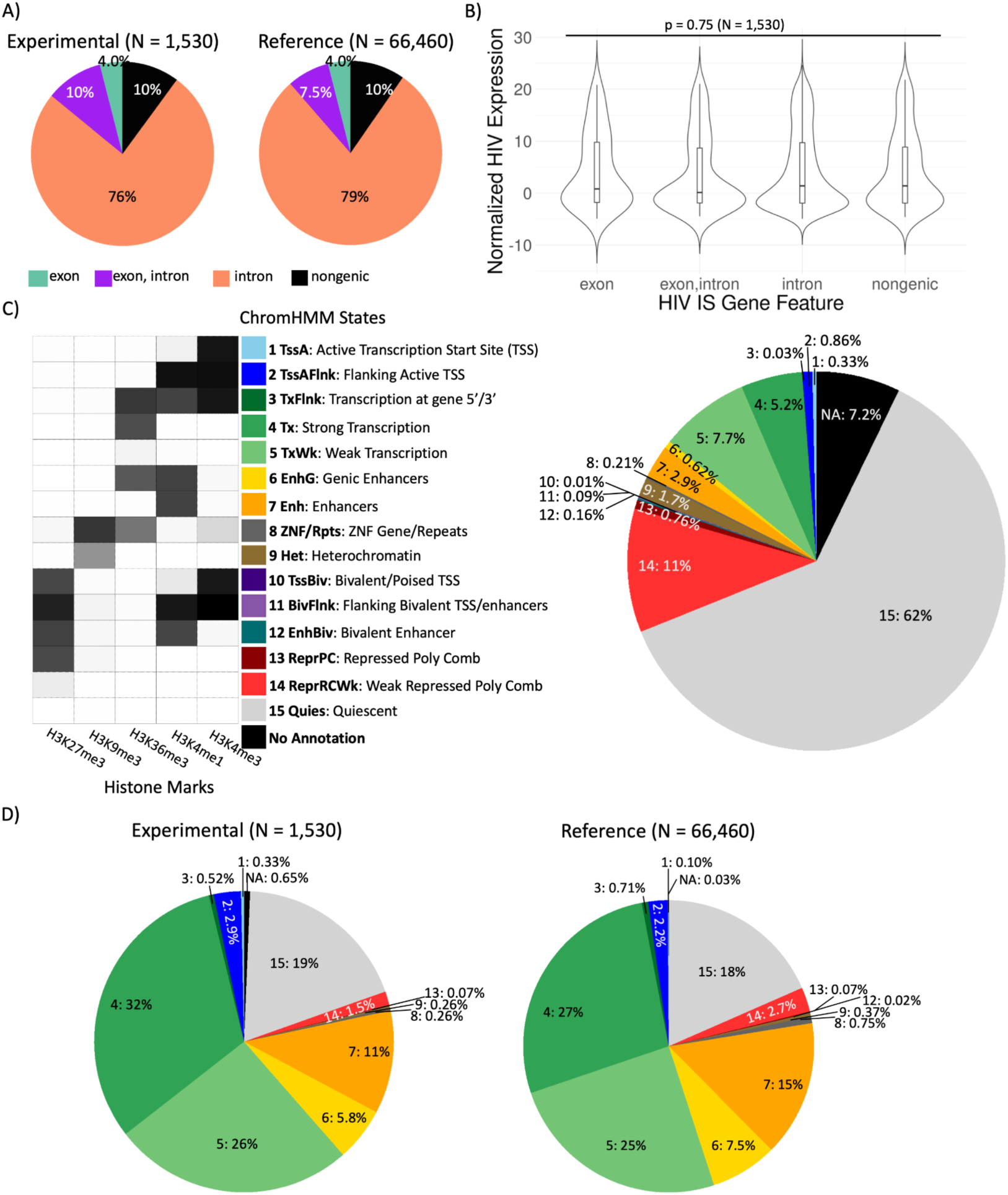
HIV integration is enriched in transcriptionally active chromatin. **A)** Pie chart showing the distribution of HIV integration sites in our dataset (“Experimental”) located within introns, exons, or intergenic regions, compared to reference data from Coffin et al. (2021). **B)** Combined boxplot and violin plot showing the distribution of HIV RNA expression in cells categorized by the genic feature surrounding the integration site. No significant difference was observed (Kruskal–Wallis test, p = 0.75; N = 1,530). **C)** Characterization of chromHMM chromatin states. The heatmap displays emission probabilities for five histone marks across the 15 chromatin states. The legend includes the state abbreviations (bold) and a short description of each state. The accompanying pie chart shows the proportion of the genome (by base pairs) assigned to each state. **D)** Pie chart showing the distribution of HIV integration sites across the 15 chromHMM-defined chromatin states, compared to the reference dataset from Coffin et al. (2021).

### HIV IS chromatin states do not influence HIV expression or accessibility

We next examined whether the chromatin state at the site of HIV integration influenced HIV expression. Overall, we found no significant correlation between HIV IS chromatin state and HIV RNA expression level (Kruskal–Wallis test, p = 0.92; **Figure 5A**). This observation held true when expression levels were analyzed using unprocessed counts and stratified by treatment condition (**Figure S5A, S5B**). However, a visual trend suggests that HIV integrated into state 9 (heterochromatin) tended to exhibit lower expression levels; this difference was not statistically significant in Kruskal-Wallis test, likely due to the small number of integration sites in this compartment (N=4). Additional data would be needed to determine whether this trend is biologically meaningful. We also assessed whether chromatin state influences the accessibility of HIV proviral DNA after integration, using the number of HIV-derived ATAC DNA fragments per cell as an approximation. Although the majority of HIV IS were located in transcriptionally active regions, there was no significant difference in HIV DNA fragment counts across chromatin states (**Figure 5B**).

**Figure 5.**
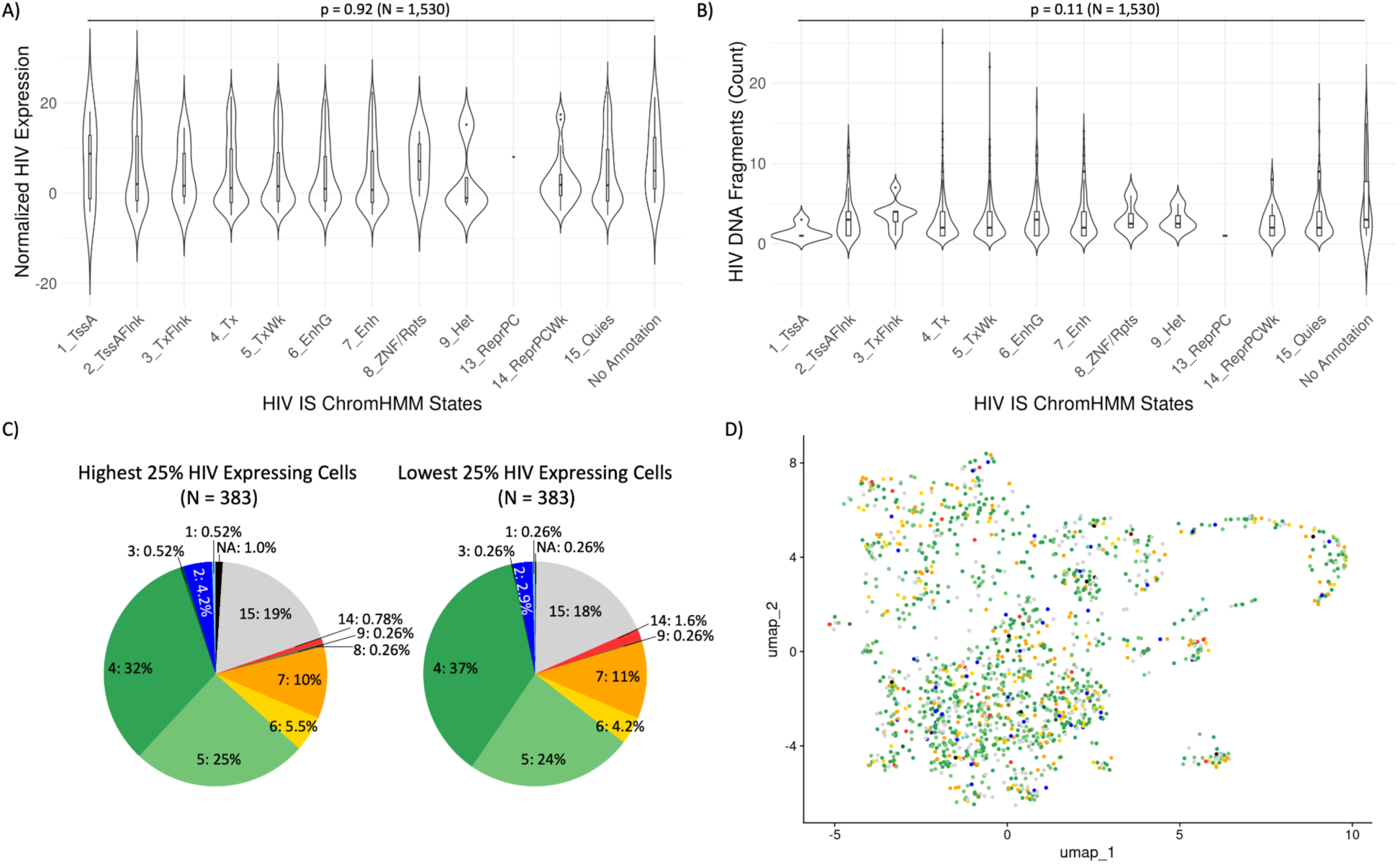
HIV IS chromatin states do not strongly influence HIV expression or accessibility. **A)** Combined boxplot and violin plot showing the distribution of HIV RNA expression across chromatin states of HIV integration sites. No significant differences were observed (Kruskal–Wallis test, p = 0.92; N = 1,530). **B)** Combined boxplot and violin plot showing the distribution of HIV DNA fragments (from ATAC-seq) per cell, categorized by HIV IS chromatin state. No significant differences were observed (Kruskal–Wallis test, p = 0.11; N = 1,530). **C)** Pie charts showing the proportion of HIV integration sites assigned to each chromatin state, stratified by the highest and lowest quartiles of HIV-expressing cells (N_total_ = 1,530; N_quartile_ = 383 per group). **D)** UMAP visualization of HIV IS+ cells colored by chromatin state, showing no strong spatial clustering of chromatin features across the transcriptomic landscape (N = 1,530).

To further investigate whether chromatin context impacts viral activity, we compared the chromatin state distribution of HIV IS from the highest and lowest quartiles of HIV- expressing cells. We observed no significant differences in integration patterns between these groups (**Figure 5C**). Additionally, mapping chromatin states onto a UMAP projection of host transcriptomes revealed no strong spatial clustering, indicating that IS chromatin context is not tightly linked to overall host gene expression profiles (**Figure 5D**). Together, these results suggest that while HIV preferentially integrates into transcriptionally active regions, the host chromatin state at the site of integration does not strongly influence HIV accessibility and expression.

### HIV integration results in insertional activation of host cell genes

After establishing that chromatin structure surrounding HIV IS does not have a strong general effect on HIV expression, we next sought to examine how the orientation of HIV integration relative to the host gene’s transcriptional direction affects viral expression. We observed that, for proviruses within host cell genes, HIV integration orientation appeared random with respect to the host gene (**Figure 6A**), and orientation had no apparent impact on HIV RNA expression (**Figure 6B**). This finding remained consistent when analyzed using unprocessed counts and stratified by treatment condition (**Figure S6**). Together with earlier results, these data suggest that HIV expression is largely independent of the genic context or orientation of its integration site.

**Figure 6.**
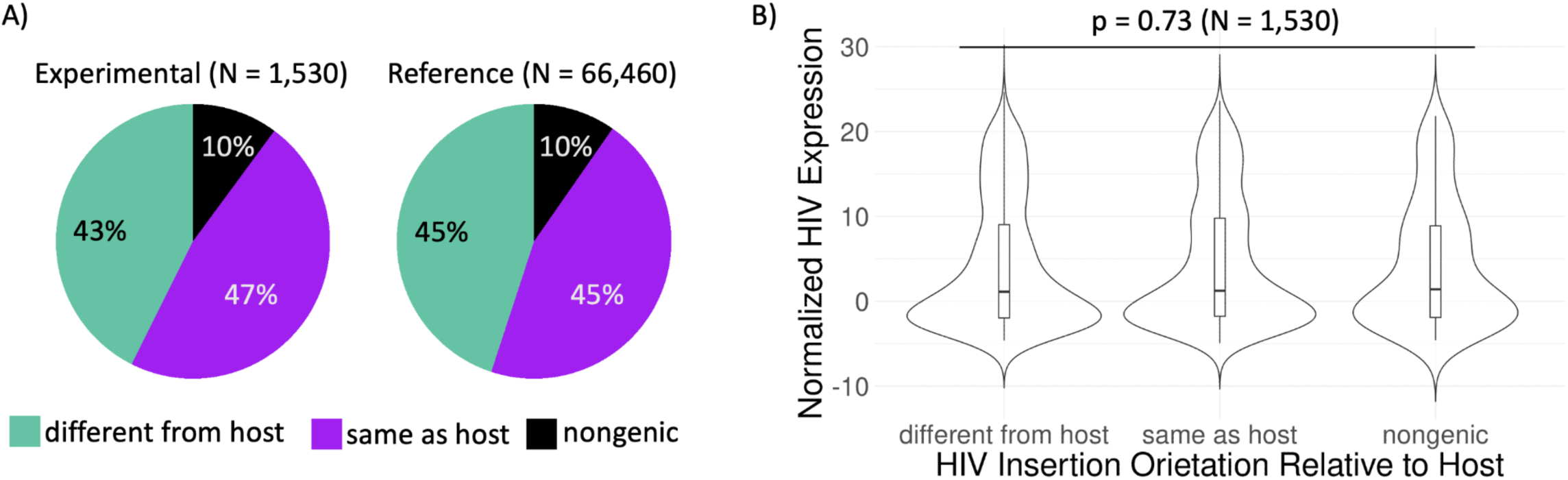
HIV insertion orientation does not influence HIV expression. **A)** Pie chart showing the proportion of HIV integration sites oriented in the same or opposite transcriptional direction as the host gene, compared to Coffin et al. (2021). **B)** Combined boxplot and violin plot showing HIV RNA expression stratified by HIV insertion orientation. No significant difference observed (Kruskal–Wallis test, p = 0.73; N = 1,530).

We next examined whether HIV affects expression of the host cell genes into which it integrates. Prior studies have described orientation-dependent insertional activation in a limited number of cases (Mellors et al. 2021), but the conditions where this effect occurs remain unclear. In our dataset, we identified individual cases in which HIV integration appeared to potently upregulate expression of the host gene, in a manner that was specific to both the orientation and location of the integration (**Figure S7**), where upregulation is defined by comparing the expression of the HIV-integrated gene in each IS+ cell to the same gene’s expression in all other cells in the dataset.

To test whether this phenomenon is generalizable, we aggregated gene expression values across HIV-integrated genes in all IS+ cells. Because Seurat normalization and integration produces expression values that are approximately normally distributed with a mean near zero and a standard deviation near one for each gene, direct comparisons across genes are valid. As a negative control, we also generated a simulated dataset randomly placed *in silico* HIV integration sites and performed the same analysis. We found that, in more than 5% of IS+ cells, expression of the integrated gene exceeded the mean expression of that gene across the population by a large margin (approximately more than 5 standard deviations than mean). This phenomenon was absent in the randomized dataset, indicating a biological effect rather than a sampling artifact. Moreover, this insertional activation occurred almost exclusively when HIV was integrated in the same orientation as the host gene (**Figure 7A**). To control for potential confounding effects from globally high gene expression cells (e.g., DE cluster 2, identified earlier in **Figure 2D**), we assessed the proportion of cluster 2 cells among insertion-activated cells. While insertion-activated cells were somewhat enriched for cluster 2 (9.7%) compared to all cells (4.9%) and IS+ cells (7.6%), the overall low proportion suggests that cluster 2 is not a major driver of the observed insertional activation (**Figure 7B**).

**Figure 7.**
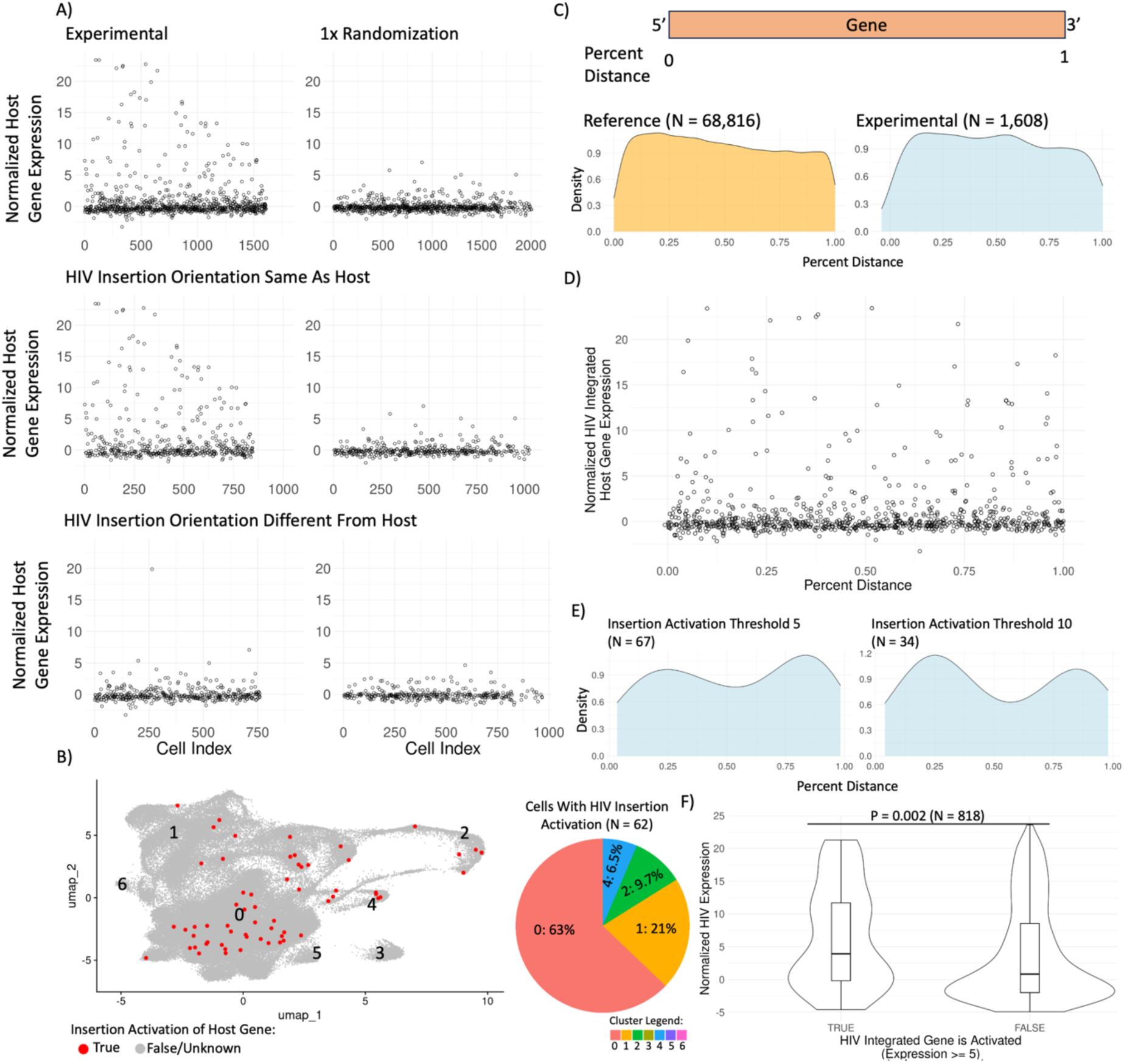
HIV integration promotes frequent insertional activation of host cell genes. **A)** Scatter plots showing the expression of HIV-integrated host genes, compared to randomly generated integration sites. Top row: all samples; middle and bottom rows: stratified by HIV insertion orientation relative to the host gene. **B)** UMAP projection of all cells, with HIV insertion–activated cells highlighted in red and the rest of the population in gray. Pie chart indicates the proportions of transcriptional clusters within the insertion-activated subset. **C)** Schematic defining percent distance of HIV IS along the gene body, with accompanying density plot comparing IS distributions in experimental dataset and Coffin et al. (2021). **D)** Scatter plot showing the relationship between insertion site location and host gene activation. **E)** Density plots showing HIV IS distribution along genes at different thresholds of host gene upregulation (e.g., approximately ≥ 5 or ≥ 10 standard deviations above the mean. **F)** Combined boxplot and violin plot showing HIV RNA expression in cells with and without host gene insertional activation. HIV expression is significantly higher in the insertion-activated group (Kruskal–Wallis test, p = 0.002; N = 818).

Next, we examined whether the position of the HIV integration site within the host gene affects the likelihood of insertional activation. HIV integrations were distributed broadly across gene bodies, with a modest enrichment near gene start sites (**Figure 7C**). While no monotonic trend emerged between percent distance along the gene and activation likelihood, density plots using increasing thresholds for defining high expression revealed bimodal enrichment: integration near the 5′ and 3′ ends of genes was more likely to be associated with extreme host gene activation (approximately 5 SD above the mean; **Figure 7D, 7E**). This enrichment diminished at higher thresholds, potentially due to reduced sample size.

In addition, we observed a weak correlation between HIV expression in IS+ cells and the expression of the host gene into which it was integrated. Notably, HIV RNA levels were significantly higher in cells where insertional activation of the host gene occurred (p = 0.002, N=818; **Figure 7F**). This suggests that insertional activation of host genes is more frequent for highly expressed proviruses.

Together, these findings suggest that HIV integration can lead to frequent insertional activation of host genes, particularly when the provirus is in the same orientation as the gene. However, the locational specificity and precise details of these events remains unclear and will require further investigation to elucidate the underlying mechanisms.

## Discussion

The influence of local genomic features surrounding HIV integration sites (HIV IS) on both viral and host gene expression remains largely uncharacterized. In this study, we used single-cell ATAC-seq to identify HIV integration sites and correlated them with host and viral transcriptional profiles obtained from matched single-cell RNA-seq data. Our findings confirm that HIV preferentially integrates into transcriptionally active regions of the genome, consistent with previous bulk analyses (Schröder et al. 2002, Craigie et al. 2012). Notably, HIV expression appeared relatively robust across diverse genomic contexts. In addition, we observed that HIV integration can upregulate the expression of host genes, particularly when the provirus is inserted in the same transcriptional orientation as the host gene.

We identified HIV integration sites using the epiVIA pipeline (Wang et al., 2020) and validated them by confirming a higher frequency of host–HIV chimeric read pairs compared to host–host interchromosomal chimeras, which likely reflect PCR recombination artifacts. Additionally, HIV reads were consistently aligned to long terminal repeats (LTRs) and oriented outward from the viral genome, a pattern suggesting genuine host–viral junctions in unsequenced parts of the DNA segment. Together, these features support the biological authenticity of the identified integration sites.

A major limitation of this approach, however, is its sensitivity. Despite successful infection being confirmed in all cells by fluorescence reporter, and ∼40% of cells containing detectable HIV DNA by ATAC-seq, we were able to confidently assign integration sites in only ∼1% of cells. This discrepancy likely reflects the low probability of capturing a host–virus junction by random pair-ended ATAC fragments. Moreover, it introduces interpretive ambiguity: the IS- population likely includes both cells lacking accessible integration sites and cells for which the integration status is inconclusive.

To address technical biases inherent to ATAC-seq, particularly the reliance on proviral accessibility (“ATAC bias”), we compared our dataset with a reference set of HIV integration sites identified via nested PCR (Coffin et al., 2021), which is independent of chromatin accessibility. We also applied Seurat integration (Hao et al. 2023) to mitigate batch effects and harmonize HIV expression across treatment conditions. After correction, small-molecule treatments had minimal impact on expression clustering, with the exception of a minor prostratin-driven cluster (Cluster 4, ∼1.2% of cells), which did not affect HIV or host gene expression. By contrast, Cluster 2 showed elevated total RNA levels and was enriched for genes associated with cell cycle phases between DNA synthesis and mitosis (**Figure S3C**). While HIV expression was elevated in this cluster regardless of integration status, Cluster 2 did not contribute significantly to insertional activation events. These findings suggest that although cell cycle–associated transcriptional activity may enhance HIV expression, it is largely independent of HIV integration site features.

Our analysis confirmed that HIV preferentially integrates into intronic regions of the genome. However, whether this preference results from an intrinsic affinity for introns or simply reflects the larger proportion of intronic sequences within genic regions remains inconclusive. Preliminary attempts to clarify this by comparing intronic and exonic base pair compositions yielded inconclusive results due to complications arising from overlapping genic annotations.

To further investigate the host genomic environment around HIV IS at a higher resolution, we applied the chromHMM model, which classifies 200 bp genomic windows based on five histone modification marks (Roadmap Epigenomics Consortium et al. 2015, Ernst et al. 2017). Our findings revealed a preferential integration of HIV into transcriptionally active chromatin states, consistent with previous reports (Craigie et al. 2012, Vasant et al. 2020). We ruled out the possibility that this integration preference results from ATAC bias by comparing our dataset with a reference dataset (Coffin et al. 2021) obtained from people with HIV. Despite differences in cell populations and sequencing methods, the proportions of HIV IS across genomic categories remained consistent, suggesting that HIV integration preferences observed are fundamental to its mechanism. However, our single-cell IS and gene expression analysis did not reveal a significant correlation between genomic features and HIV viral expression. Several factors may explain this observation. First, as mentioned earlier, ATAC-seq bias enriches for transcriptionally active and accessible sites, although we found that IS+ cells still exihibited a wide range of HIV expression levels and a large fraction of cells without detectable viral RNA expression. Second, while visual differences in HIV expression were observed among different genomic categories, there is no statistical significance, possibly due to small sample sizes in some chromatin states (e.g., heterochromatin, where only four cells were identified). A more efficient IS identification method could improve statistical power and validate these trends. Third, for higher-resolution chromHMM states, it is possible that while HIV preferentially integrates into accessible regions, the chromatin state of the surrounding genome does not necessarily dictate the provirus’s transcriptional accessibility. Given that chromHMM assigns chromatin states based on 200 bp windows and HIV is approximately 10 kb long, integration could alter or override the local chromatin landscape. Finally, HIV expression may be inherently robust to the local chromatin environment around the integration site, with previous reports of silencing in transcriptionally inactive regions representing outlying cases (Jiang et al., 2020).

Recent reports have demonstrated that intact proviruses are preferentially preserved in high repetitive chromatin regions during long term ART and in elite controllers (Lian et al. 2021, Einkauf et al. 2022). While we observed a modest trend towards lower expression in H3K9me3 rich chromatin compartments, this effect was not significant, possibly due to the low numbers of integration sites within our dataset. Nevertheless, our data indicate that for the large majority of HIV integration sites, expression is relatively robust to the surrounding chromatin configuration, indicating that HIV transcription mechanisms typically override the influence of local chromatin. Our findings should, however, be considered in light of the caveat that our integration site identification approach is somewhat biased for cells with active viral gene expression, due to its reliance on the presence of viral ATACseq reads. Also, our results do not rule out the influence of integration site on HIV expression – studies of HIV integration sites in clonal T cells lines indicate diverse and stable patterns of expression for identical proviruses with different sites and orientations (Winslow et al. 1993, Jordan et al. 2001, Jordan et al. 2003, Han et al. 2008), arguing that integration site details have an important impact on HIV expression. However, we speculate that this effect may be due to precise details of integration that have not yet been accounted for, rather than simply the host chromatin state at the integration site.

Previous studies have suggested that HIV proviruses integrated into ZNF family genes, centromeric, or telomeric regions exhibit downregulated expression that promotes viral latency (Einkauf et al. 2022, Huang et al. 2021, Lian et al. 2021, Jiang et al. 2020). However, exploratory analysis of this dataset did not reveal significant differences in HIV expression between proviruses integrated into ZNF genes and those integrated elsewhere, likely due to the limited sample size of ZNF-integrated cells (n = 15 of 1,530 total IS+ cells). Notably, all ZNF-integrated proviruses were found exclusively in LRA-treated samples (iBET, prostratin, and SAHA), suggesting potential treatment-induced detection bias for ZNF integrated provirus. Similarly, only one cell with centromeric integration (on chromosome 11) was identified, containing a single HIV RNA copy with normalized HIV expression value of 0.27. No telomeric integrations were detected. The low discovery rate for integration sites in these regions may be attributed to ATAC sequencing bias toward accessible chromatin or mapping issues within repetitive centromeric and telomeric regions. Therefore, while the single centromeric case agrees with previous findings of reduced expression, the regulatory effects of integration into ZNF genes, centromeric, and telomeric regions on HIV expression remain inconclusive in this dataset due to limited sample sizes and sequencing method biases.

Finally, we also observed that HIV integration can frequently activate the expression of host genes. These insertional activation events appear to depend on the orientation of the provirus relative to the host gene and may also be influenced by the specific location of the integration site within the gene body. While orientation-dependent activation was clearly observed, since it occurred almost exclusively when the provirus and host gene were aligned, we could not conclusively establish location specificity. This may reflect limitations in our current analytic approach, which uses percent distance along the gene as a positional indicator. A more precise definition involving regulatory elements could help resolve this question in future studies. Future experiment could also focus on determining the causal correlation between HIV integration orientation and position and host gene expression.

In conclusion, our study provides key insights into the genomic landscape of HIV integration and its effects on host and viral transcription. We confirm that HIV preferentially integrates into transcriptionally active genic regions—a pattern that appears to be an intrinsic viral preference across diverse conditions. However, we did not observe a strong correlation between integration site features and viral expression, likely due to the intrinsic robustness of HIV transcription. Notably, our results suggest that HIV integration can induce host gene upregulation in an orientation-dependent manner. Future work combining large datasets, refined regulatory mapping, and functional assays will be critical to fully elucidate how integration site characteristics influence HIV latency and reservoir dynamics.

## Acknowledgements

This work was supported by the following grants from the National institutes of Health: NIAID #5-R01AI143381 and NIAID #5-UM1AI16456. The funders had no role in study design, data collection and analysis, decision to publish, or preparation of the manuscript. We thank the UNC HIV cure center, UNC Center for AIDS Research (P30-AI-050410), and the UNC Flow Cytometry Core Facility.

## Methods

### Dataset acquisition and alignment of scATACseq and scRNAseq data

The datasets analyzed in this study were obtained from the Gene Expression Omnibus (GEO) database under accession number GSE242997, and some data were previously published in Manickam et al. 2024. CD4+ T cells were isolated, infected with a GFP-encoding strain of HIV (HIV-dreGFP/Thy1.2), and, for some samples, treated with the indicated compounds, following the protocol described in Peterson et al. 2023. Combined single-cell ATAC and RNA sequencing were performed using the 10x Genomics multiomic platform (CG000365.RevB) (Manickam et al. 2024), and libraries were sequenced on an Illumina NextSeq platform (CG000338.RevD) (Manickam et al. 2024). Prepared libraries underwent paired-end sequencing on an Illumina NextSeq platform. Sequences were aligned using the 10x Genomics Cell Ranger ARC software. A custom human-HIV reference genome, incorporating the HIV-dGFP/Thy1.2 construct, was built with the mkref function (Manickam et al. 2024). Alignment against this custom reference genome was performed using the cellranger-arc count function. For subsequent analyses, the .bam files generated from this alignment were used for DNA information, while the filtered_feature_bc_matrix files were used for gene transcriptome analysis.

### Identification of HIV integration site with epiVIA

Paired chimeric reads were identified in the .bam files from the alignment, defined as reads with two ends aligned to different chromosomes (in the alignment .bam file, these reads have a 7th column value that is not an “=” sign). The baseline paired chimeric rate was calculated by dividing the number of paired chimeric reads on a chromosome by the total number of reads aligned to that chromosome. HIV integration sites were identified following the epiVIA pipeline described by Wang et al. 2020. The .bam files from scATAC-seq alignments served as inputs. Custom reference genome FASTA files, incorporating the human-HIV and HIV-dGFP/Thy1.2 sequences (Manickam et al. 2024), were indexed using BWA (Li et al. 2009) and used for integration site mapping. In cases where a cell had multiple HIV IS, the whole cell was excluded from the study.

### Annotation of local chromatin structure by chromHMM

The chromHMM annotations, based on primary T helper cells from peripheral blood (ID: E043) from the NIH Roadmap Epigenomics Consortium, were used to define chromatin states (Roadmap Epigenomics Consortium et al. 2015, Ernst et al. 2017). These annotations were generated using a 15-state Hidden Markov Model trained on five core histone marks (H3K4me3, H3K4me1, H3K36me3, H3K27me3, and H3K9me3) and provided as the E043_15_coreMarks_hg38lift_mnemonics_sorted.bed file (Roadmap Epigenomics Consortium et al. 2015, Ernst et al. 2017). HIV integration sites identified using epiVIA were processed and matched to these chromatin annotations. The GenomicRanges package in R was used to assign a chromatin state to each HIV integration site (Lawrence et al. 2013, R Core Team 2023). In cases where an HIV IS had two chromHMM states (because the HIV IS coordinates may be a range), a random one was retained.

### Annotation of the orientation of HIV provirus relative to host transcriptional direction

HIV-integrated genes were re-annotated using the main 23 chromosomes from GENCODE V46 via the UCSC Table Browser (Mudge et al. 2025, Perez et al. 2015, Karolchik et al. 2004), with gene symbols extracted from hg38.kgXref. The orientation of each HIV integration site, as determined by epiVIA, was compared to the orientation of the gene it integrated into and classified as either “same as host,” “different from host,” or NA when HIV was not integrated in a gene.

### Annotation of gene features around HIV integration site

The GENCODE V46 reference described previously (Perez et al. 2015, Karolchik et al. 2004) was used to define gene features. BED files for introns and exons were generated directly using the UCSC Table Browser (Mudge et al. 2025, Perez et al. 2015, Karolchik et al. 2004). Non-genic regions were created by subtracting intron and exon intervals from the whole genome BED file using bedtools subtract function (Quinlan et al. 2010). With the GenomicRanges package in R (Lawrence et al. 2013, R Core Team 2023) each HIV integration site was assigned a feature type “exon,” “intron,” or “nongenic,” based on the interval containing it. If an HIV integration site was both intronic and exonic due to overlapping genes, it was categorized as “exon,intron.”

### Reference dataset of HIV integration sites

The reference dataset of HIV integration sites was obtained from the publicly accessible Retrovirus Integration Database v2.0 (Coffin et al. 2021). It was lifted to the hg38 genome assembly using the hg19-to-hg38 chain file (Perez et al. 2025, Hinrichs et al 2006) and annotated with the GENCODE V46 reference for compatibility with the experimental data (Mudge et al. 2025). All integration features were annotated using the methods described above.

### Normalization and integration of single-cell gene expression data

The gene expression data were processed using the Seurat software suite (Hao et al. 2023). Doublets were identified and removed from each of the 16 samples using scDblFinder in R (Germain et al. 2022). The data were then normalized using Seurat’s SCTransform pipeline with default settings, following the “Using SCTransform in Seurat” tutorial (Hao et al. 2023). After normalization, the 16 samples were integrated following the “Introduction to scRNA-seq Integration” tutorial (Hao et al. 2023).

### Differential expression clustering of single cells based on gene expression data

A copy of the gene expression data, independent from the previous section, was made. After doublets were removed as described previously, HIV expression was removed from the expression matrix. Normalization and integration followed the same pipeline as previous. The data were clustered using Seurat’s clustering pipeline, following the “Guided Clustering Tutorial” (Hao et al. 2023). The biological implications of clusters were assessed by browsing the list of significantly differentially expressed genes (log2 fold change ≥ 5) in Enrichr (Chen et al 2013, Kuleshov et al. 2016, Xie et al. 2021).

### Simulation of random HIV integration sites and integrated genes

To assess the influence of HIV integration events on host gene expression, randomly generated sites were compared with experimentally derived HIV integration sites. These random sites were distributed proportionally across intronic, exonic, and non-genic regions to match the observed distribution in the actual dataset. Each random site was assigned a cell barcode and HIV orientation corresponding to the feature type (e.g., pseudo-exon sites were matched to the barcodes and orientations of real exon sites). No barcodes were reused in the 1:1 randomization process, ensuring an unbiased comparison. Randomization was performed using a fixed seed for reproducibility. The simulated sites were annotated with associated genes using the GENCODE V46 reference as described in previous sections (Mudge et al. 2025, Perez et al. 2015, Karolchik et al. 2004). Gene expression values for the corresponding cell barcodes were extracted from the actual Seurat object.

### Definition of HIV integration site percent distance in a gene

The relative position of an HIV integration site within a gene was measured using percent distance, where the start and end of the gene were defined as 0 and 1, respectively. The percent distance for each integration site was calculated using the following formula:

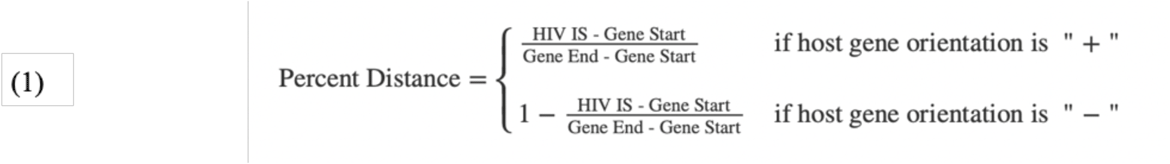

### Statistical analysis

The statistical tests in this study were conducted in R (R Core Team 2023). For comparisons among more than two groups, a Kruskal–Wallis test was used to identify differences (R Core Team 2023), followed by a post-hoc Dunn’s pairwise test when appropriate (Dinno 2024). For features with two groups, a Wilcoxon rank-sum test was performed (R Core Team 2023). Correlations were calculated using the Spearman method (R Core Team 2023). All graphics were generated using base R, ggplot2, or matplotlib in Python (R Core Team 2023, Wickham 2016, Hunter 2007).

### Code and data availability

All code, data, and figures used in this analysis are available in the following Google Drive directory: https://drive.google.com/drive/folders/1oGw428eWQ24q-YlfzZUveIP7tFZzUipJ?usp=share_link

## Glossary

ART: Antiretroviral Therapy
DE: Differential Expression
HIV IS: HIV Integration Sites
LRA: Latency Reversal Agents
LTR: Long Terminal Repeats
PWH: People With HIV

---

**Supplemental Table 1.** Sample information. All donors were seronegative for HIV.

| GEO | Sample | Cell Count | Cell Source | Treatment |
| --- | --- | --- | --- | --- |
| GSE242997 | unstimulated1 | 16,474 | Donor 1 | DMSO (0.05%, 24hr) |
|  | unstimulated2 | 8,101 | Donor 2 | DMSO (0.05%, 24hr) |
|  | iBET1 | 16,285 | Donor 1 | iBET151 (75nM, 24hr) |
|  | iBET2 | 8,180 | Donor 2 | iBET151 (75nM, 24hr) |
|  | Prostratin1 | 15,215 | Donor 1 | Prostratin (75nM, 24hr) |
|  | Prostratin2 | 9,473 | Donor 2 | Prostratin (75nM, 24hr) |
|  | SAHA1 | 15,852 | Donor 1 | SAHA (500nM, 24hr) |
|  | SAHA2 | 6,168 | Donor 2 | SAHA (500nM, 24hr) |
| Unpublished | unstimulated3 | 3,353 | Donor 3 | Methanol (0.01%, 0hr) |
|  | unstimulated4 | 1,713 | Donor 4 | Methanol (0.01%, 0hr) |
|  | unstimulated5 | 1,441 | Donor 5 | Methanol (0.01%, 0hr) |
|  | THC3hr1 | 4,980 | Donor 3 | THC (500nM, 3hr) |
|  | THC6hr1 | 3,702 | Donor 3 | THC (500nM, 6hr) |
|  | THC6hr2 | 1,466 | Donor 4 | THC (500nM, 6hr) |
|  | THC6hr3 | 2,084 | Donor 5 | THC (500nM, 6hr) |
|  | THC12hr1 | 3,127 | Donor 3 | THC (500nM, 12hr) |

**Supplemental Figure 1.**
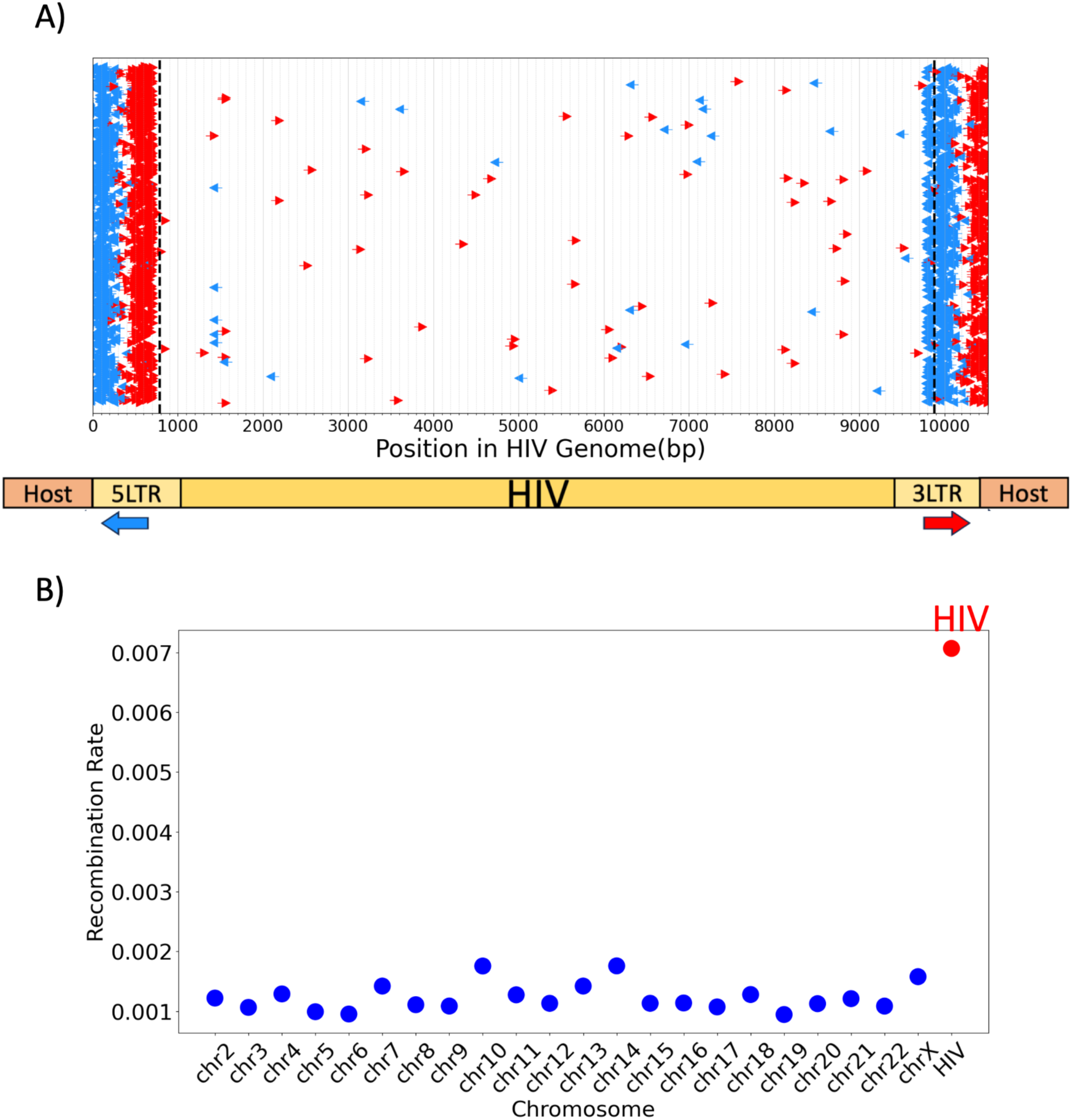
Characterization of HIV integration sites (IS). **A)** Map of the HIV genome. Each arrow represents a paired chimeric DNA fragment aligned to the HIV genome, with arrow direction and color indicating read orientation. The long terminal repeats (LTRs) are marked by dotted lines. Due to the identical sequences between the 5′ and 3′ LTRs, reads mapping to these regions cannot be distinguished by the aligner and are assigned randomly. However, paired-end sequencing favors short fragments, suggesting that reads mapping deeper into an LTR likely originate from the opposite end of the provirus. This is illustrated in the schematic at the bottom of (A). **B)** Pair chimeric rate across chromosomes. The pair chimeric rate is defined as the fraction of chimeric reads relative to total reads for a given chromosome. Blue points represent host–host pair chimerics (one read mapping to one host chromosome, the other to a different host chromosome), while red points represent host–HIV pair chimerics.

**Supplemental Figure 2.**
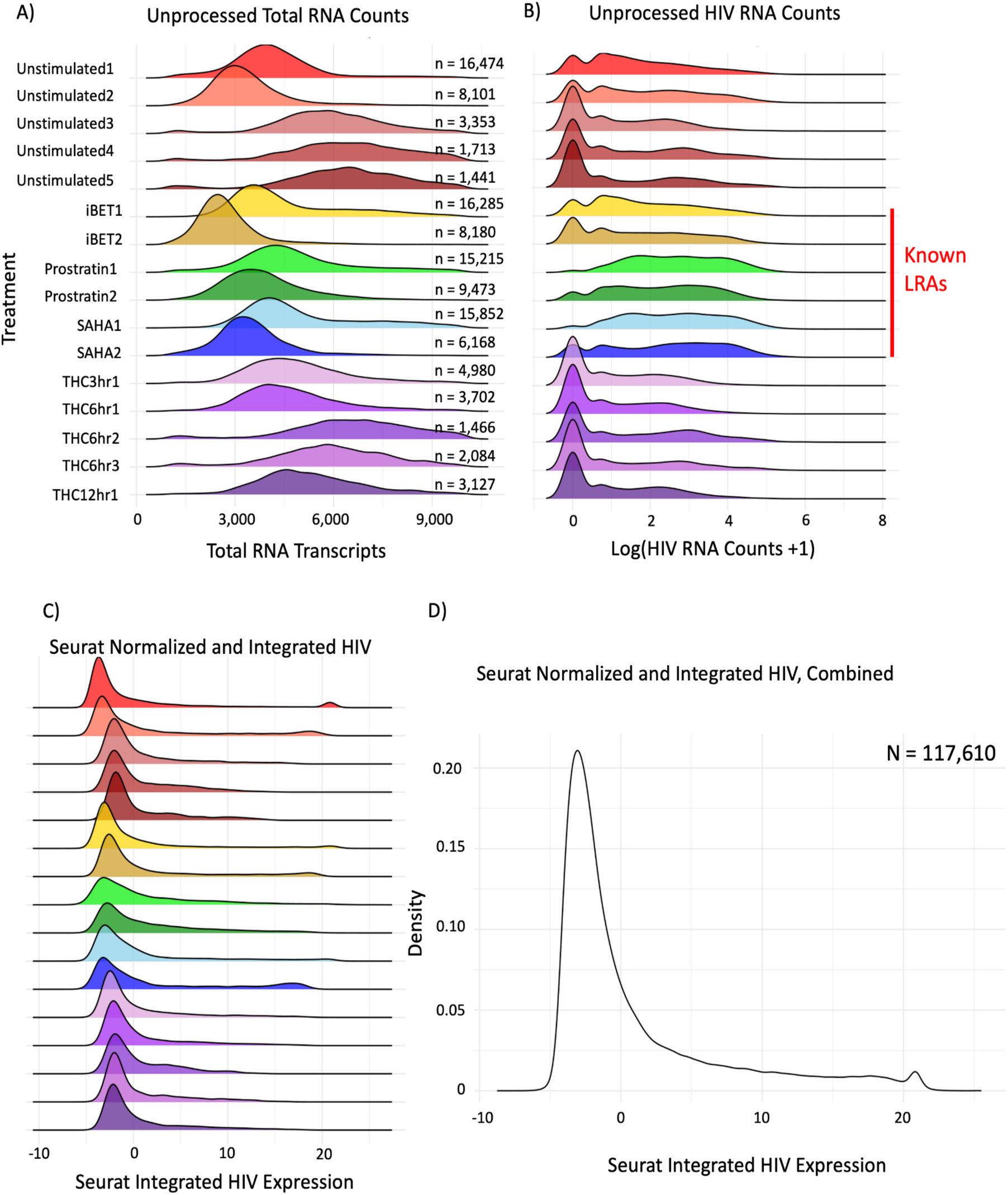
HIV mRNA distribution. **A)** Density plot of total RNA counts per cell, stratified by treatment. **B)** Density plot of unprocessed HIV mRNA counts (logged for visualization). **C)** Density plot of HIV mRNA expression after SCT normalization and Seurat integration, stratified by treatment. No units are shown, as Seurat integration alters the scale and interpretation of expression values. **D)** Density plot of aggregated SCT-normalized and Seurat-integrated HIV mRNA expression across all cells (N = 117,610).

**Supplemental Figure 3.**
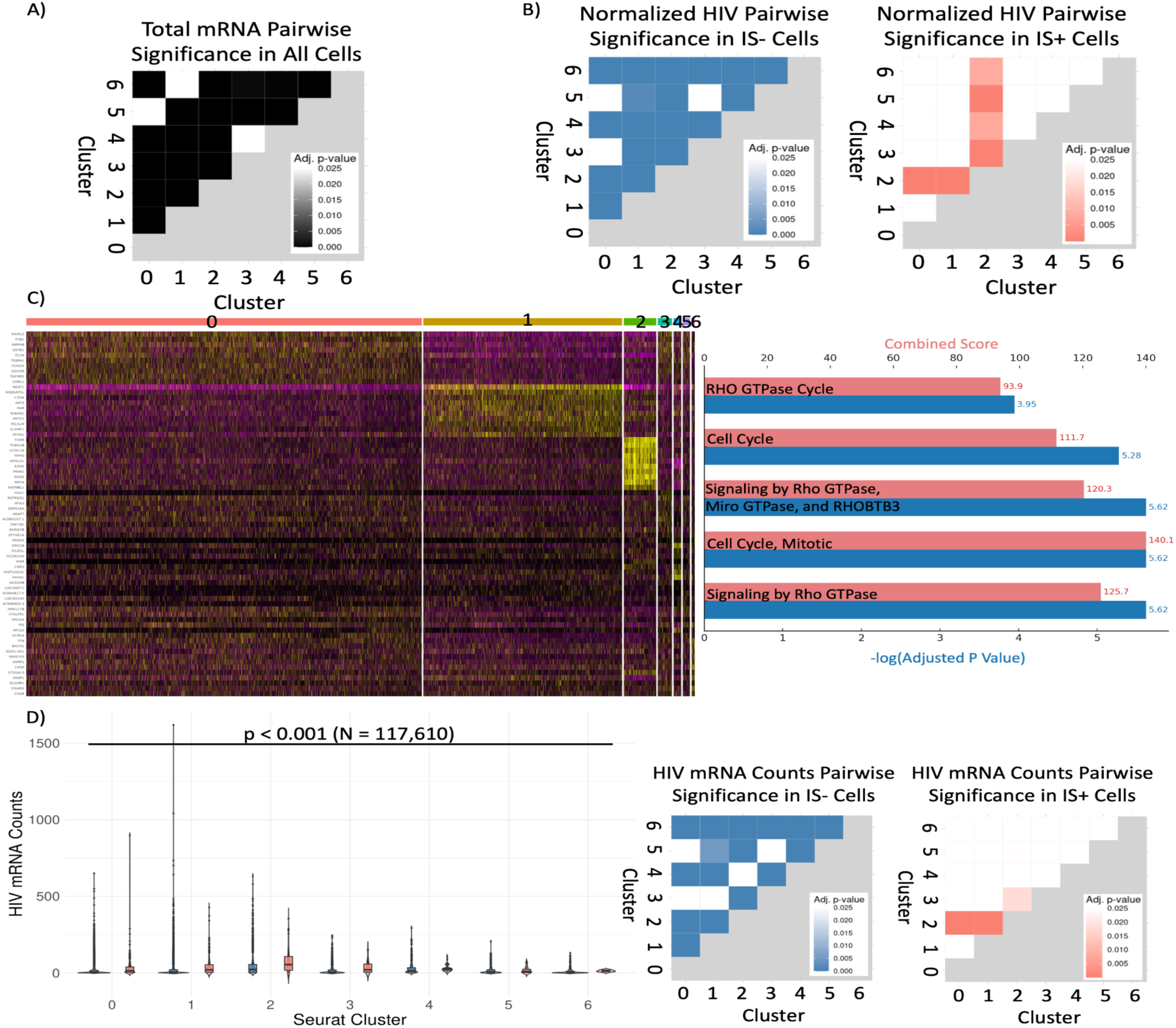
Transcriptomic characterization of differential expression clusters. **A)** Heatmap of p-values from pairwise comparisons of total mRNA content per cell across clusters, following the overall significant result shown in Figure 2D. Color intensity indicates statistical significance (Dunn’s test). Due to large sample sizes and imbalance between cluster sizes, Dunn’s test may identify statistically significant but biologically trivial differences. Visual inspection of Figure 2D suggests Cluster 2 as the most distinct. **B)** Heatmaps of p-values from pairwise comparisons of normalized HIV expression per cell across clusters, following the overall significance reported in Figure 2E. **C)** Heatmap displaying the top 10 differentially expressed genes for each cluster. The right panel shows top 5 ontology analysis results for Cluster 2 using Enrichr (Chen et al 2013, Kuleshov et al. 2016, Xie et al. 2021). A comprehensive list of DE genes per cluster is provided in the Data Availability section. **D)** Boxplot and violin plot showing unprocessed HIV mRNA counts per cell across clusters, used to validate the trends observed in normalized data (Figure 2E). Kruskal–Wallis tests confirm significant differences across clusters for both IS- and IS+ populations (p < 0.001 for both). Blue indicates IS- cells; orange indicates IS+ cells.

**Supplemental Figure 4.**
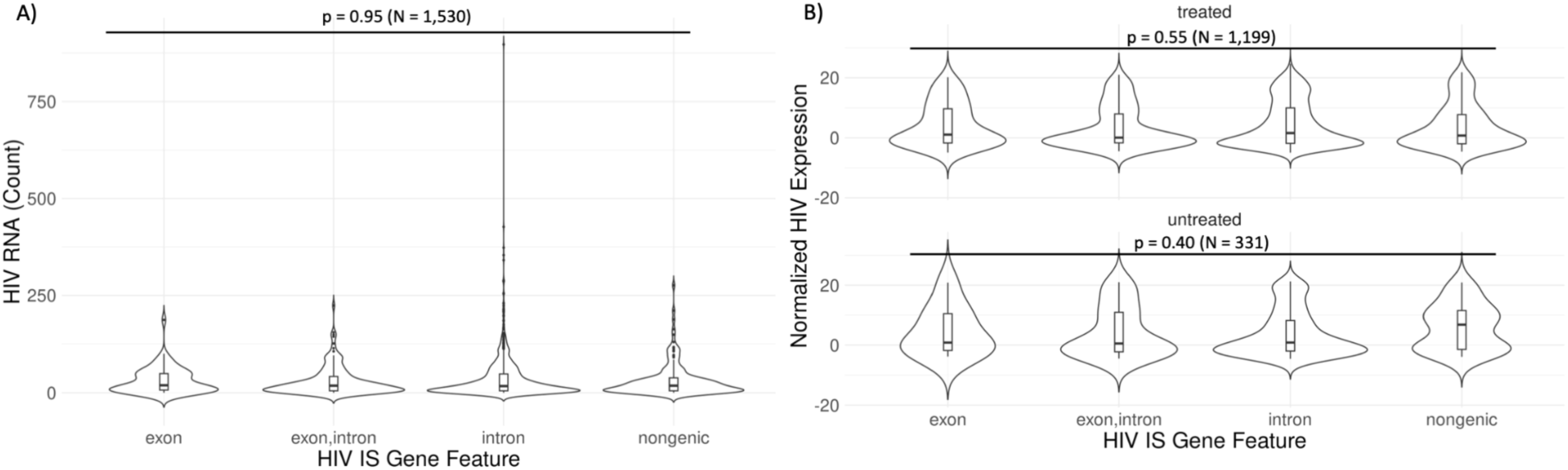
Sample treatment does not influence HIV expression across genic context. **A)** Replication of Figure 4B using unprocessed HIV RNA counts. Combined boxplot and violin plot showing the distribution of unprocessed HIV expression levels, categorized by the genic feature (intron, exon, or intergenic) surrounding the HIV integration site. No significant differences were observed (Kruskal– Wallis test, p = 0.95; N = 1,530). **B)** Replication of Figure 4B stratified by small molecule treatment. Combined boxplot and violin plot showing normalized HIV RNA expression by genic feature, separated by treatment condition. No significant differences were observed in either group (p_treated_=0.55, N_treated_ = 1,199; (p_untreated_=0.40, N_untreated_ = 331).

**Supplemental Figure 5.**
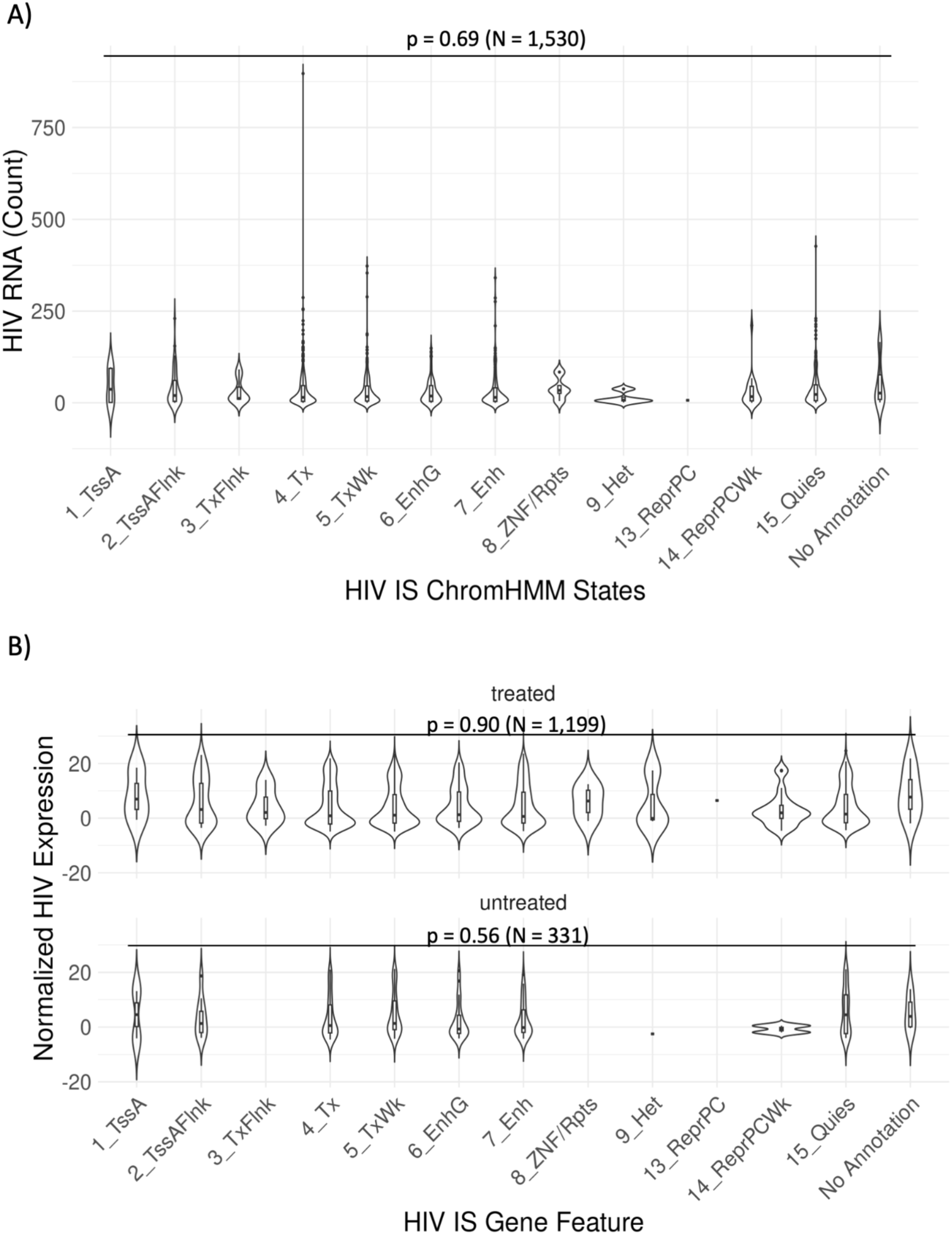
Sample treatment does not influence HIV expression across chromatin states. **A)** Replication of Figure 5A using unprocessed HIV RNA counts. Combined boxplot and violin plot showing HIV expression across chromatin states. No significant differences were detected (p=0.69, N = 1,530). **B)** Replication of Figure 5A stratified by small molecule treatment. Combined boxplot and violin plot showing normalized HIV RNA expression across chromatin states in treated and untreated groups. No significant differences were observed in either condition (p_treated_=0.90, N_treated_ = 1,199; p_untreated_=0.56, N_untreated_ = 331)

**Supplemental Figure 6.**
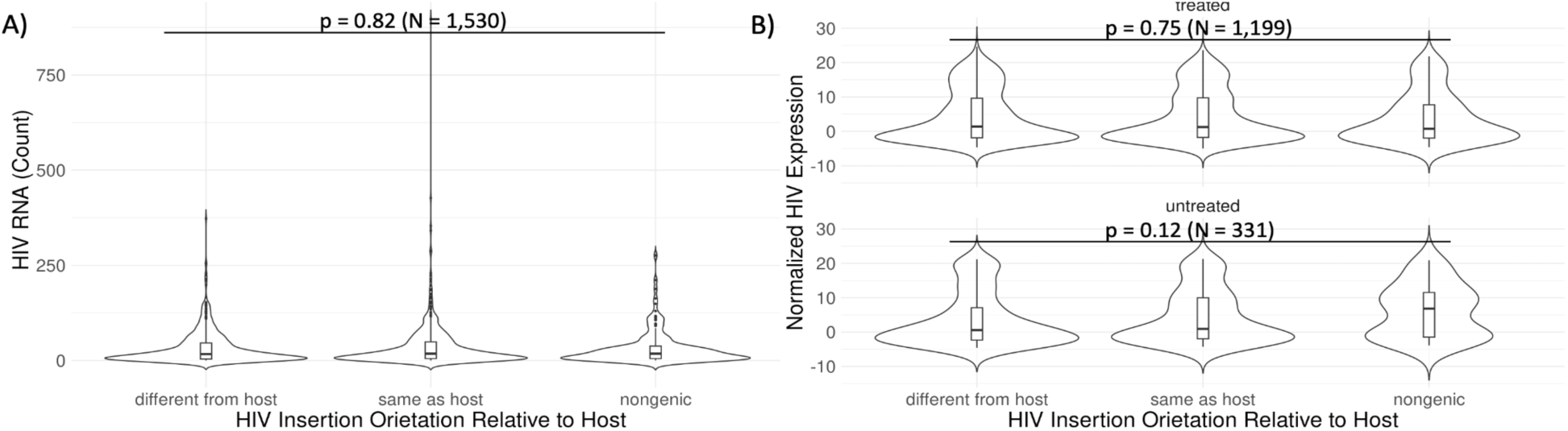
Sample treatment does not influence HIV expression across insertion orientations relative to the host gene. **A)** Replication of Figure 6B using unprocessed HIV RNA counts. No significant difference observed across orientations (Kruskal–Wallis test, p = 0.82; N = 1,530). **B)** Replication of Figure 6B stratified by small molecule treatment. Kruskal-Wallis test not significant for either group (p_treated_=0.75, N_treated_ = 1,199; p_untreated_=0.12, N_untreated_ = 331).

**Supplemental Figure 7.**
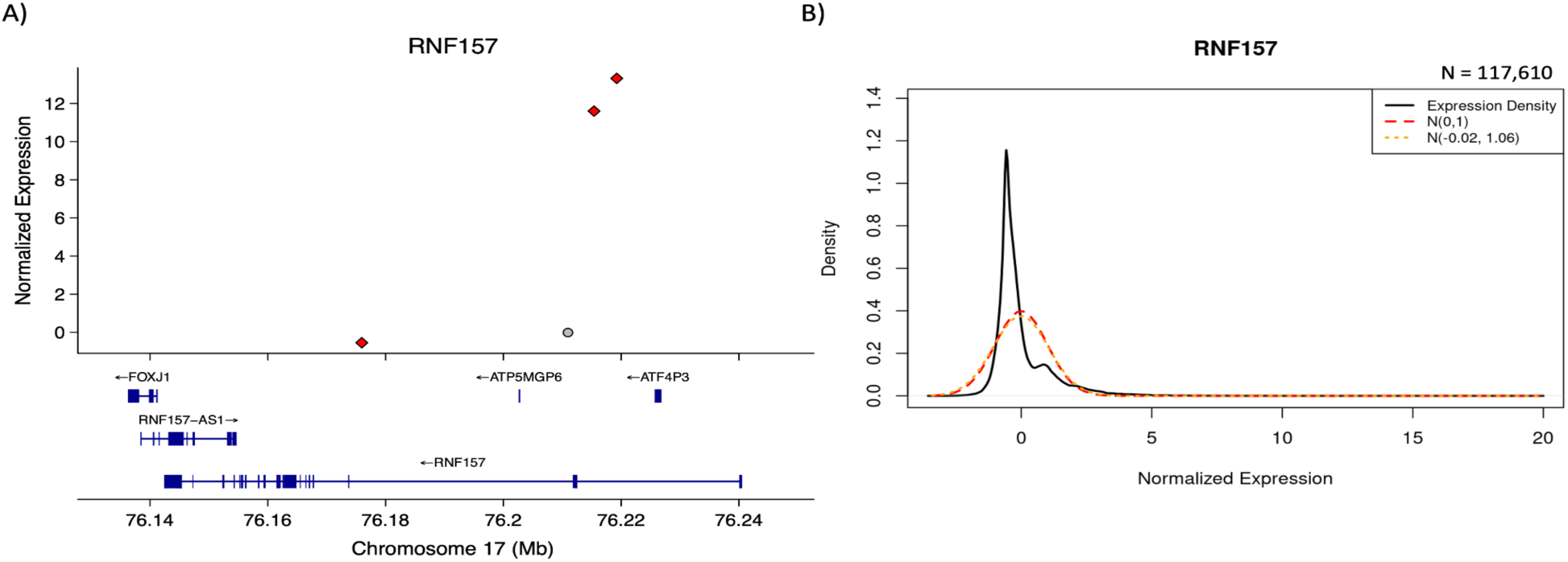
Case study shows HIV insertion activation is specific to HIV IS location and orientation relative to the host gene. **A)** Scatter plot showing RNF157 expression in individual cells with HIV integrated into the RNF157. Each point represents a cell; red diamonds indicate HIV in the same transcriptional orientation as RNF157, and gray dots indicate the opposite orientation. **B)** Density plot of RNF157 expression across all 117,610 cells after Seurat normalization. The black curve represents the empirical distribution; red and orange curves represent reference normal distributions (mean = 0, SD = 1; and RNF157-specific parameters, respectively). Additional case studies for all 65 insertion-activated genes are available in the Data Availability section.

**Supplemental Figure 8.**
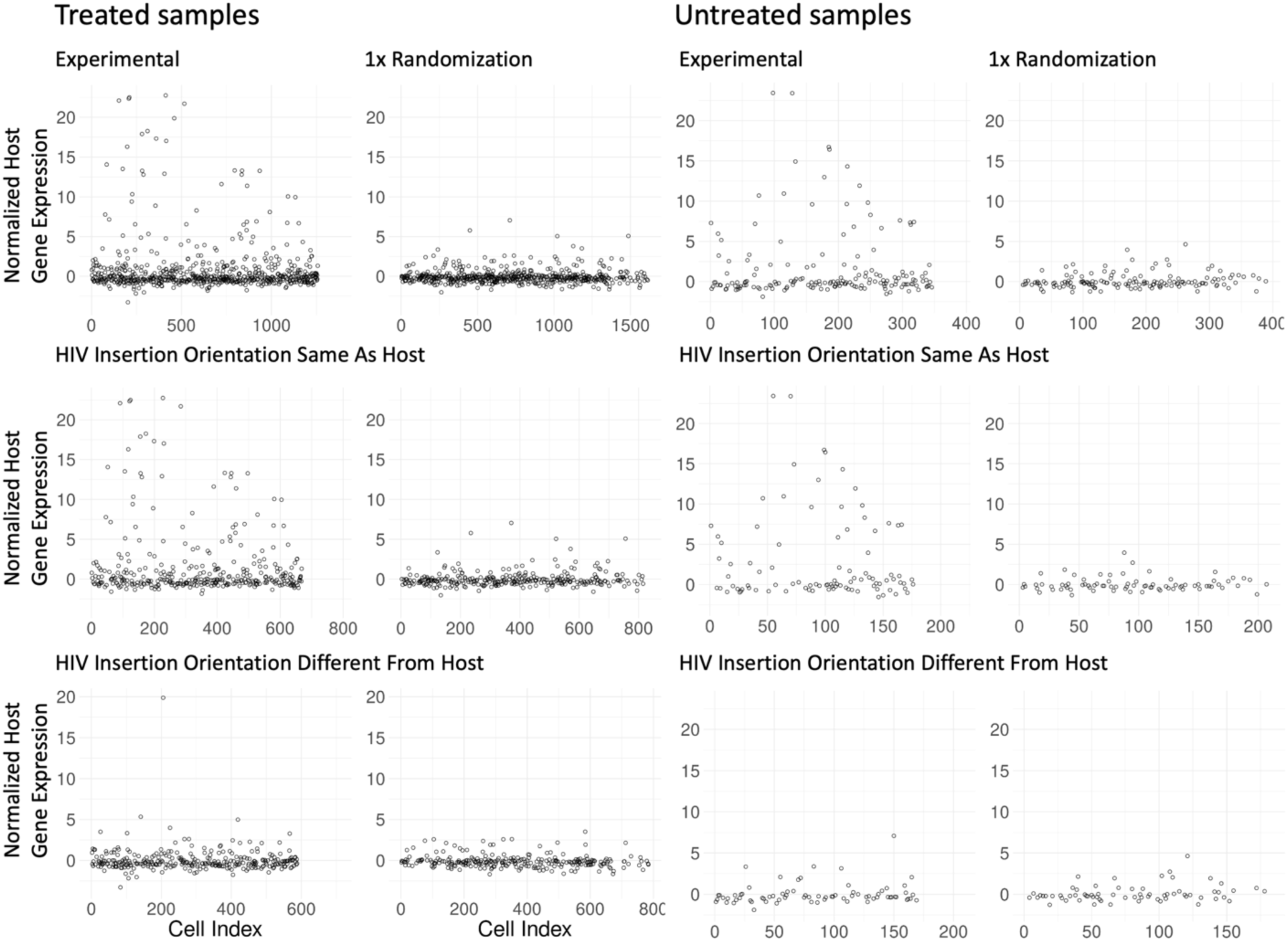
HIV insertion activation of host gene expression is independent of treatment. Further stratification of Figure 7A by small molecule treatment condition. Insertional activation remains consistent across treatment groups.

